# A Fully Automated, Faster Noise Reduction Approach to Increasing the Analytical Capability of Chemical Imaging for Digital Histopathology

**DOI:** 10.1101/425835

**Authors:** Soumyajit Gupta, Shachi Mittal, Andre Kajdacsy-Balla, Rohit Bhargava, Chandrajit Bajaj

## Abstract

High dimensional data, for example from infrared spectral imaging, involves an inherent trade-off in the acquisition time and quality of spatial-spectral data. Minimum Noise Fraction (MNF) developed by Green *et al*. [1] has been extensively studied as an algorithm for noise removal in HSI (Hyper-Spectral Imaging) data. However, there is a speed-accuracy trade-off in the process of manually deciding the relevant bands in the MNF space, which by current methods could become a person month time for analyzing an entire TMA (Tissue Micro Array). We propose three approaches termed ‘Fast MNF’, ‘Approx MNF’ and ‘Rand MNF’ where the computational time of the algorithm is reduced, as well as the entire process of band selection is fully automated. This automated approach is shown to perform at the same level of reconstruction accuracy as MNF with large speedup factors, resulting in the same task to be accomplished in hours. The different approximations of the algorithm, show the reconstruction accuracy vs storage (50*×*) and runtime speed (60*×*) trade-off. We apply the approach for automating the denoising of different tissue histology samples, in which the accuracy of classification (differentiating between the different histologic and pathologic classes) strongly depends on the SNR (signal to noise ratio) of recovered data. Therefore, we also compare the effect of the proposed denoising algorithms on classification accuracy. Since denoising HSI data is done without any ground truth, we also use a metric that assesses the quality of denoising in the image domain between the noisy and denoised image in absence of ground truth.

## Introduction

Chemical imaging is an emerging technology in which every pixel or voxel of an image contains hyperspectral data, often consisting of hundreds or thousands of data points. The spectrum at each pixel resolves the chemical components at that point and, thus, provides the molecular profile of the sample [2–4]. Computer algorithms that can process the data to information useful for a particular problem often require a specific data quality, at that spectral resolution, that often determines scanning (signal averaging) time. In addition to the chemical signature of the data, another benefit of these technologies is that workflows can be automated with fully digital analysis of the data [5–7]. For example, Fourier Transform Infrared (FT-IR) spectroscopic imaging is emerging as an automated alternative to human examination in studying disease development and progression by using statistical pattern recognition [8–13]. For a practical protocol for tissue imaging, as demonstrated in at least one instance of tissue histopathology, the signal-to-noise ratio (SNR) of 4*cm^−1^* resolution spectral data needs to be more than 1000 : 1 [12]. To achieve this SNR, especially for the emerging high definition IR imaging [14–16], extensive signal averaging is required. The need for signal averaging increases acquisition time (SNR *~ 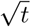*), in turn, increasing acquisition time [17] to the extent that clinical translation becomes impractical. Signal processing approaches to reduce noise has previously been suggested to mitigate this crippling increase in integration time by mathematical methods to utilize correlations in data to reduce noise but suffer from two major drawbacks. First, given the large size of the data, the mathematical operations require computer processing often comparable to the acquisition time itself [18]. Second, such methods invariably try to separate data into informative and noisy components; subsequently, a manual selection step is required to identify the information-bearing components thus compromising the automation benefits of using spectroscopic imaging for tissue analysis [19].

One class of mathematical transform techniques for noise reduction utilize the property that noise is uncorrelated whereas spectra (signals) have a high degree of correlation. In a transform domain, hence, the signal becomes largely confined to a few eigenvalues whereas the noise is spread across all. Noise reduction can be achieved by retaining eigenvalue images that correspond to high signal content and computing the inverse transform. All the eigenvalue data contain signal and noise but the relative proportion of the signal to noise which forms a threshold criterion for inclusion of specific eigenimages in the inverse transform. Inclusion of too many will not allow for significant noise rejection, while inclusion of too few would result in loss of fine spectral features. Hence, identifying eigenvalues corresponding to high signal content is an important step in the noise reduction process.

One widely used algorithm was provided by Green *et al*. [1] that applies Minimum Noise Fraction (MNF) to order spectral components in terms of SNR in the transformed space. It assumes that the covariance for the raw data ∑_*Y*_ and the noise ∑_δ_ can both be estimated. Similar to principal component analysis (PCA) that orders the components in terms of variance, after transformation in MNF space, the top components are chosen and filtered and rest are zeroed out. This reduced basis is then used for inverse transformation into the signal space. Noise Adjusted Principal Components (NAPC) transform by Lee *et al*. [20] is a reformulation of the MNF transform in terms of noise whitening process. While MNF and NAPC transforms are mathematically equivalent, the latter consists of a sequence of two principal component transforms: First to whiten the data (de-correlate noise from data); Second to perform eigen decomposition on the modified covariance matrix, to order the underlying data by SNR.

Our goals in this work are to address the major challenges in noise rejection using mathematical methods. Specifically, first, we aim to provide criteria for unsupervised band selection, thereby maintaining objectivity of the data analysis protocol and reducing analysis time within the data processing pipeline by dispensing with the need for manual intervention. Second, we aim to re-examine the mathematical formulation of the MNF approach to speed up the process of computation of the forward and inverse transform MNF vectors. Specifically, we examine two novel approaches: (a) use of a truncated singular value decomposition (SVD) variant that is computationally more efficient, and (b) the use of a randomized variant of the above that is also memory efficient and has a reconstruction accuracy-memory trade-off depending on the application. Finally, we seek to improve the signal processing methods to provide a higher confidence in the consistency and accuracy of the noise rejection pipeline. We propose a comparison between acquired and denoised biomedical images using a robust metric, thereby providing better denoising guarantees in terms of both root mean square error (RMSE) and structural similarity.

## Materials and methods

### Sample Preparation and Data Collection

A paraffin embedded breast tissue microarray (*BR*1003) consisting of 101 cores were obtained from US Biomax, Inc. The unstained sections of the TMA were placed on a *BaF*_2_ salt plate for IR imaging. The sections were deparaffinized using a 24*h* hexane bath. High Definition data was acquired using the Agilent Stingray imaging system with 0.62 numerical aperture and a 128 *×* 128 focal plane array. A spectral resolution of 4*cm^−1^* along with a pixel size of 1.1*µm* was obtained at the sample plane. The final FTIR sample has a spatial dimension of 11620 *×* 11620 pixels and spectral dimension of 1506 channels.

### Classification

A random forest classifier was used to differentiate between the different histologic classes of a tissue sample. Labeled pixels for each class were obtained by the cases annotated by a pathologist. In this study, we have used a four class model separating benign epithelium from malignant epithelium. Finally, to assess the performance of the classifier, sensitivity and specificity is calculated for all the classes to generate the receiver operating characteristic curve. The area under this curve signifies the diagnostic potential of the model.

## Methods

In practical situations, noiseless data *d_j_* is recorded by instruments as a noisy signal estimate *y_j_*, due to fluctuations in detector current, backgrounds and source/instrument factors, which is modeled as:

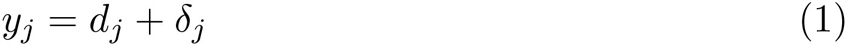

where *y_j_* ∈ ℝ^*S*^ is the actual data collected by the apparatus and *δ_j_* ∈ ℝ^*S*^ is the noise in the same pixel. The goal is to estimate *δ_j_* given *y_j_*, so we can best estimate the true signal *d_j_* such that the relevant features (peak positions, peak heights, relative peak spacing etc.) in the spectrum are preserved.

Given a HSI *Y_orig_* ∈ ℝ^*W*×H×S^, where *W, H* are the spatial dimensions (width and height respectively), it has been restructured into a 2*D* matrix *Y ∈* ℝ^*N*×S^, where *N* = (*W × H*) is the number of pixels, *S* is the number of spectral channels. Each column *y_i_, ∀i* = [1 : *S*] represents the reshaped image for the *i^th^* spectral band and each row *y_j_, ∀j* = [1 : *N*] represents the spectral signature for the *j^th^* pixel. The *i^th^* entry of the spectral vector *y_j_* ∈ ℝ^*S*^ of each pixel *y_ij_* determines the absorbance value of the tissue at that wavenumber. Wavenumbers are defined as inverse of wavelength and have units *cm^−1^*. The recorded value at each wavenumber has unit *absorbance/au*, where *au* is arbitrary unit.

### Mathematical Background

Assuming additive noise only, raw data can be represented as *Y* = *D* + *δ*, where *Y* = *{Y*_1_, …, *Y_S_}* and *D* and δ are the uncorrelated signal (actual spectral data with baseline included) and noise components of the raw data *Y*. *Cov{Y}* = Σ_*Y*_ = Σ_*D*_ + Σ*_δ_*, where Σ_*D*_ and Σ_*δ*_ are the covariance matrices of *D* and *δ* respectively. Noise Fraction (NF) for the *i^th^* band is defined as the ratio of noise variance to the total variance for that band. Similarly Signal to Noise ratio (SNR) for the *i^th^* band is defined as the ratio of signal variance to the noise variance for that band.

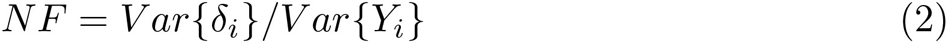

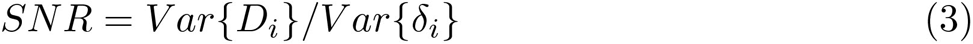

MNF is the set of linear transformation (*Y_MNF_*)_*i*_ = *Y ϕ_i_*, for *i* = 1, …, *S*, such that the SNR for (*Y_MNF_*)_*i*_ is maximum among all linear transformations orthogonal to (*Y_MNF_*)_*j*_, for *j* = *i* + 1, …, *S*. All the transformation vectors in MNF space follow 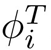, *∀i* = [1 : *S*]. Maximization of the noise fraction leads to a numbering of bands that gives decreasing image quality with increasing component number. The SNR for (*Y_MNF_*)_*i*_ in MNF space can be formulated:

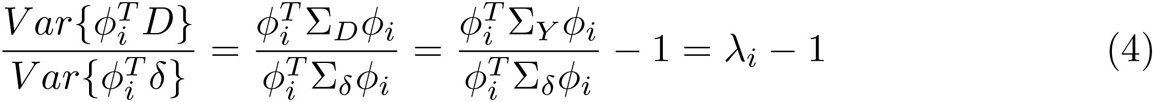

The Noise fraction itself can then be re-factored as follows:

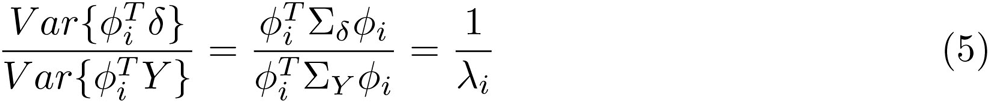

The vectors *ϕ_i_* are thus the real, symmetric eigenvectors of the eigenvalue problem:

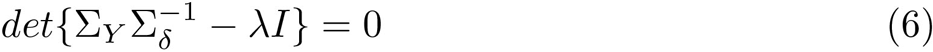

Hence *ϕ_i_* are the eigenvectors of ∑_Y_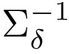, and *λ_i_*, eigenvalue corresponding to *ϕ_i_*, equals to the noise fraction in (*Y_MNF_*)_*i*_. Also, λ_1_ ≥ λ_2_ *≥…≥ λ_S_*, so that the components show decreasing image quality. Thus SNR is given by *λ_i_* − 1.

### Geometric Interpretation

As the data *D* and noise *δ* are additive and uncorrelated, we can write:

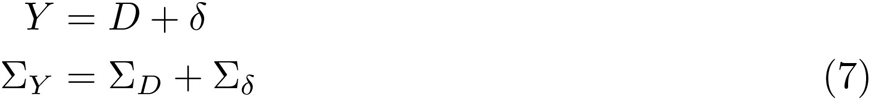

Let the spectral decomposition of Σ_*δ*_ be Σ_*δ*_ = *E*Λ_*δ*_*E^T^*. Rotating Σ_*Y*_ in Eq.7 with eigenvector matrix *E* and rescaling it using the inverse-square root of the noise singular valuesΛ_*δ*_, results in new covariance matrix where the contribution of noise component has been turned into an identity matrix.

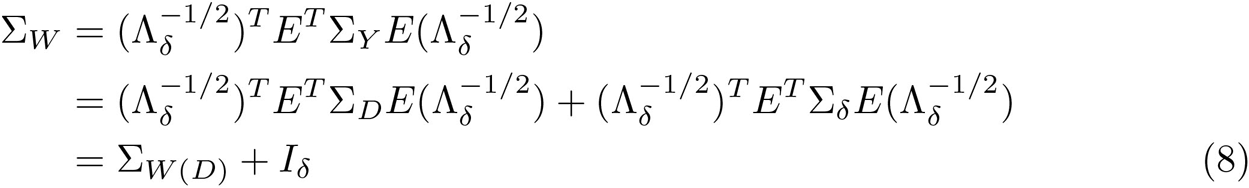

Let the eigen decomposition of Σ_*W*_ be Σ_*W*_ = *G*Λ_*MNF*_ G^T^. Rotating Σ_*W*_ in Eq.8 with eigenvector matrix *G* results in:

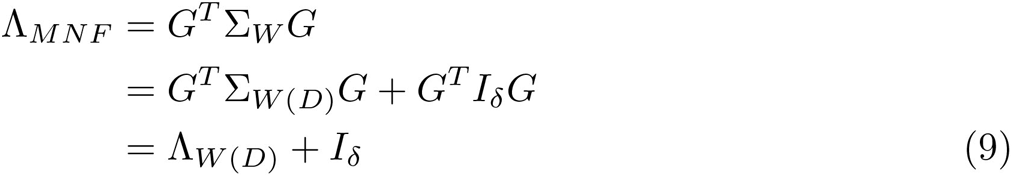

The series of transforms that we used to change the original data covariance matrix Σ_*Y*_ into Λ_*MNF*_ is given by the transformation vector 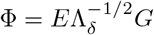 which we call the MNF projection vectors and _*MNF*_ estimates of SNR of the data.

### Optimizing MNF

Expanding the entire MNF transform, we can infer the following:

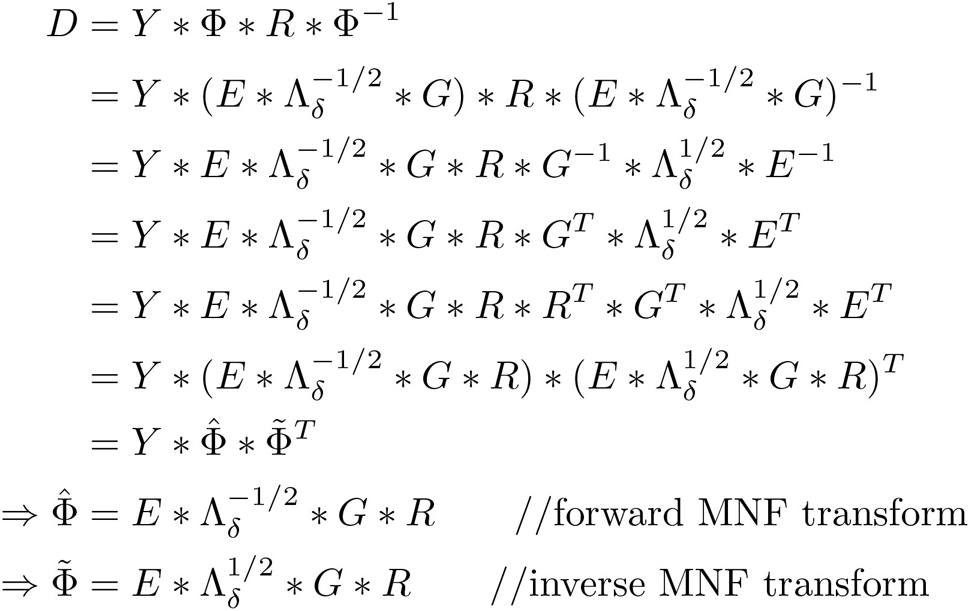

Since *R* is a block identity matrix *R* = 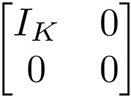, introducing an extra *R^T^* term keeps the value of the expression unaltered as *R * R^T^* = *R*. Writing the MNF transform in this way, we ensure that we skip the costly matrix inversions of the MNF vectors. Also *G * R* is effectively choosing the top *K* eigenvalues of *G*. Therefore we can reduce the computation cost by finding the reduced rank-*K* SVD of Σ_*Y*_. The Λ_*δ*_ matrix inversion can also be replaced by more efficient versions due to its diagonal structure.

### Automatic Band Selection

The optimal value of *K* can be determined by inspecting the entries of Λ_*MNF*_ = *SNR* + 1 which is a diagonal matrix. The Rose criteria [21] states that an SNR of at least 5.0 is needed to be able to distinguish image features at 100% certainty. We select the top *K* bands in the MNF space for which *SNR* = Λ_*MNF*_ −1 ≥ 5.0. Automating this process is the main computational speed factor that brings down the processing time from days down to few hours.

### Fast MNF

By exploiting the MNF formulation (see Supplemental Material), we can avoid all inverse operations and replace them with transpose, thereby making computations faster. Owing to the symmetric structure of covariance matrices, we also compute the singular value decomposition using eigen decomposition which is faster. Also, the transformation matrices are of size (*S × K*) instead of (*S × S*) where *K ≪ S*. This is the main factor responsible for the algorithmic speedup.

### Approx MNF

Since *K ≪ S*, it is inefficient to compute the full spectral decomposition of the covariance matrix. Empirically, it was observed over different datasets, that the optimal value of the automatically selected *K* is 2 − 3% of the total number of bands *S*. Hence we compute only a rank *K*̂ truncated SVD of the whitened covariance matrix. This results in reduced computation time as well as memory. The standard solutions to truncated SVD include the power iteration algorithm and the Krylov subspace methods. Since power iteration is unstable at times due to the structure of the singular values, we use a version of the Block Lanczos method [22]. We set *K*̂ = 0.03 *× S* and compute the rank *K*̂-SVD, then let the band selection criteria to decide the optimal *K*.

### Rand MNF

Although the block Lanczos algorithm can attain machine precision, it inevitably goes many passes through Σ_*W*_, and it is thus slow when Σ_*W*_ is large or does not fit in memory. To circumvent this scenario, we use a faster randomized and memory efficient version which computes the *K*̂-SVD of Σ_*W*_ up to 1 + *∊* Frobenius norm relative error [23].

### Error Metric

In the absence of ground truth images of denoised data, we use a non-reference image quality metric this is simple and easy to use. The Method Noise Image (MNI) [24] metric aims at maximizing the structure similarity between the input noisy image and the estimated image noise around homogeneous regions and the structure similarity between the input noisy image and the denoised image around highly-structured regions, and is computed as the linear correlation coefficient of the two corresponding structure similarity maps.

## Results

### Setup

For all the experimentation, a standalone machine with Intel Xeon *E5* − 1660@3.20GHz CPU and 64GB of RAM was used. Software for the simulations, results and plots include Matlab and ENVI.

### Complexity Analysis

Table 1 shows the algorithmic time and space complexity in terms of memory usage for the different MNF versions. The best algorithm in terms of time-space-accuracy is Approx MNF. This is because it computes the best rank-K SVD with little loss in its approximation or memory usage. Depending on how much one wants a memory-accuracy trade-off, one may choose to switch to use Rand MNF, as it is a randomized version with approximation error guided by its parameters. For a typical FTIR spectrum with *S ~* 1500 bands, the algorithm estimated the optimal number of bands to be *K ~* 30, resulting in a efficiency factor of 50*×*.

**Table 1.** Comparison of Time and Space complexity of MNF versions. *O*: big-O complexity. *nnz*(*X*): # non-zero elements in *X*. *N*: # pixels, *S*: # spectral bands and *K*: # chosen bands.

| Algorithm | Time | Space |
| --- | --- | --- |
| 1. MNF | $\mathcal{O}(S^3 + NS^2)$ | $\mathcal{O}(NS + S^2)$ |
| 2. Fast MNF | $\mathcal{O}(S^3 + NSK)$ | $\mathcal{O}(NK + S^2)$ |
| 3. Approx MNF | $\mathcal{O}(S^2K + NSK)$ | $\mathcal{O}(NK + SK)$ |
| 4. Rand MNF | $\mathcal{O}(nnz(\Sigma_W)K + NSK)$ | $\mathcal{O}(NK + SK)$ |

### Timing Analysis

Standard MNF is worst in terms of time complexity as it computes the full eigen decomposition of Σ_*W*_ (𝒪(*S*^3^)) and uses all the *S* eigenvectors to transform the input data into MNF space (𝒪(*NS*^2^)). In comparison, Fast MNF still computes the full decomposition of Σ_*W*_ (𝒪(*S*^3^)), but uses the Band selection criteria to retain only the top *K* eigenvalues. Hence only the top *K* eigenvectors are used to transform the data into MNF space (𝒪(*NSK*)). For Approx MNF, we only compute a truncated rank *K*̂-SVD of Σ_*W*_, since it was empirically observed that the top *K* chosen bands lie within *K*̂ = 2 *–* 3% of *S*, thereby highly reducing the computational cost (𝒪(*S*^2^*K*)). Again, only the top *K* eigenvectors are used to transform the data into MNF space (𝒪(*NSK*)). Rand MNF uses a randomized algorithm to compute a truncated rank *K*̂-SVD of Σ_*W*_, hence it is much more efficient in terms of speed (𝒪(*nnz*(Σ_*W*_))*K*), as the values in Σ_*W*_ will only be non-zero for channels which are correlated in terms of signal. Only the top *K* eigenvectors are used to transform the data into MNF space (𝒪(*NSK*)). Refer to Fig.1 for runtime. In all the three variants with *S ~* 1500 and *K ~* 30, the algorithmic scaling of *SK* instead of *S*^2^ reduces runtime by *~* 50*×*. For the BR1003 data, the entire MNF denoising process was reduced from *~* 1 month to *~* 2 hours. This clearly shows that the larger the data size, the better is the scaling of the proposed algorithms, thereby massively reducing the computational time of denoising for large TMA and associated datasets.

**Fig 1.**
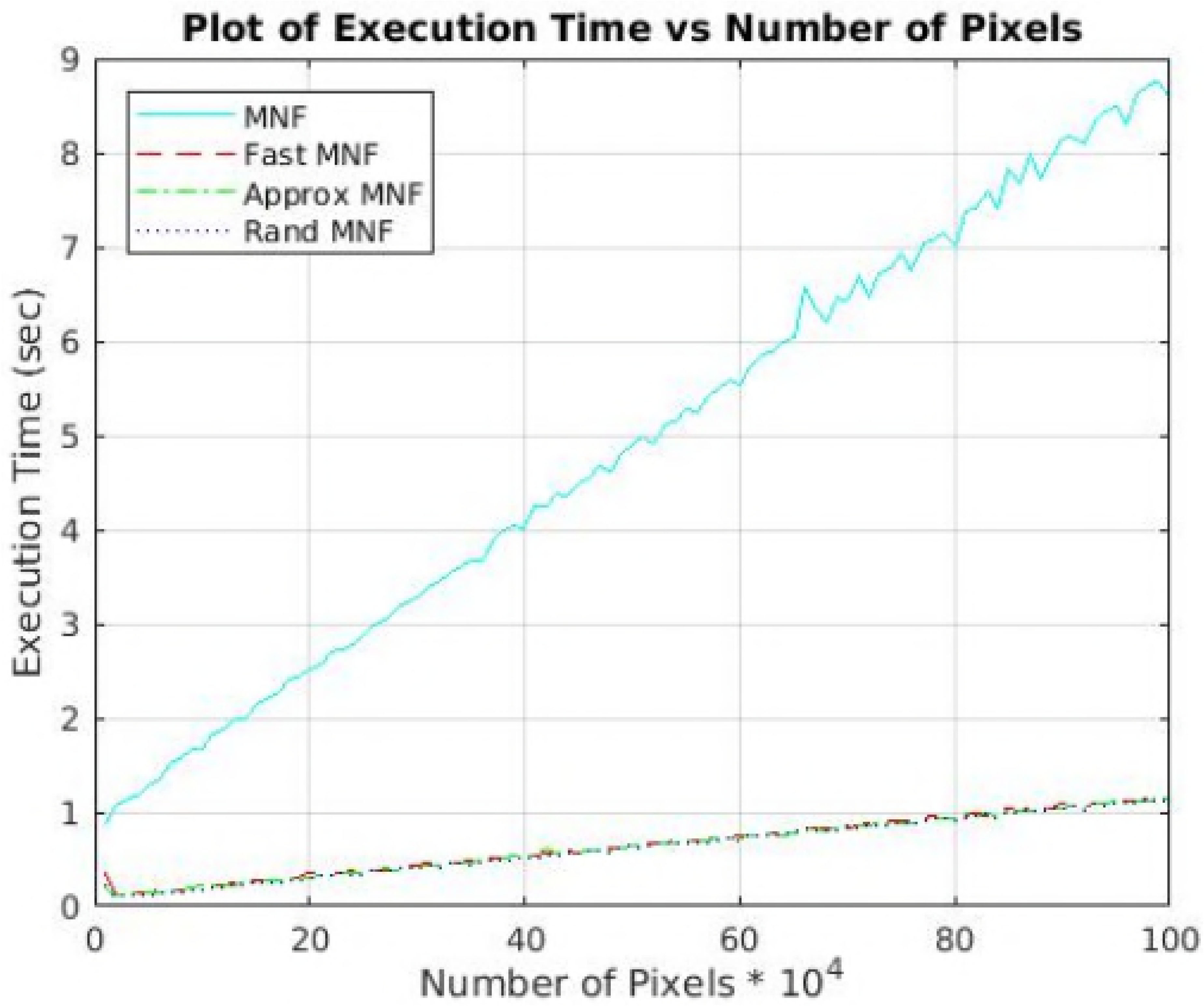
Runtime comparison of different implementations of MNF. The figure illustrates how time scales with varying size of the input data for proposed versions and optimizations of the standard MNF. The size is varied from 10*K* pixels (100 *×* 100) to 1000*K* pixels (1000 *×* 1000). Compared to the standard MNF, all the other versions have a speedup factor of *~* 10. This speedup is obtained by utilizing the fact that we do not need all the forward MNF vectors, but only the top *K* ones that are digitally calculated, avoiding manual selection. Also, a set of optimized matrix operations have been implemented, for instance replacing all inverse operations with transpose for improved computational performance. Due to BLAS optimized modules in Matlab, matrix multiplications are efficiently distributed across the cores automatically. Note: The standard MNF runtime includes automatic band selection.

**Note:** Since the structure of the noise is unknown, we cannot instinctively perform the rank-*K* approximation of Σ_δ_, because depending on experiment and instrument there is no guarantee on the strength of noise present in the raw data. For this reason, in the first step of eigen decomposition, we choose to retain bands which accounts for 99% variance in the noise bands. This further reduces the computational time of the process (*~* 3*×* extra speedup).

### Space Analysis

Standard MNF computes the full SVD decomposition (𝒪(*S*^2^)). The transformed data in MNF space contains all *S* bands (𝒪(*NS*)). Fast MNF again does the full SVD decomposition (𝒪(*S*^2^)). However the data in transformed domain only contain *K* bands (𝒪(*NK*)). Both Approx MNF and Rand MNF compute the truncated rank *K* decomposition, hence there are *K* eigenvectors each of *S* dimension (𝒪(*SK*)). The data in the transformed domain contains only *K* bands (𝒪(*NK*)). For the FTIR data with *S ~* 1500 bands and *K ~* 30, we achieve a RAM space saving of *~* 50*×*, allowing us to process more data simultaneously in one go.

### Denoising Profiles

The improvements offered by the different versions of the MNF presented in this study are illustrated in Fig. 2. The extent of denoising both in the spectral and spatial domain is approximately the same for all the different MNF algorithms. Fig. 2.A. depicts the spatial detail offered by different MNF versions with zoomed in sections in Fig. 2.C. and Fig. 2.B. shows the horizontal signal profile across the sample. Next, spectral profiles are compared across the different algorithms with reference spectrum (without MNF) to illustrate the extent of noise removal in each case (Fig. 2.D.). It can be seen that even with a speed up factor of *~* 10 there is no significant reduction in the spectral and spatial image quality.

**Fig 2.**
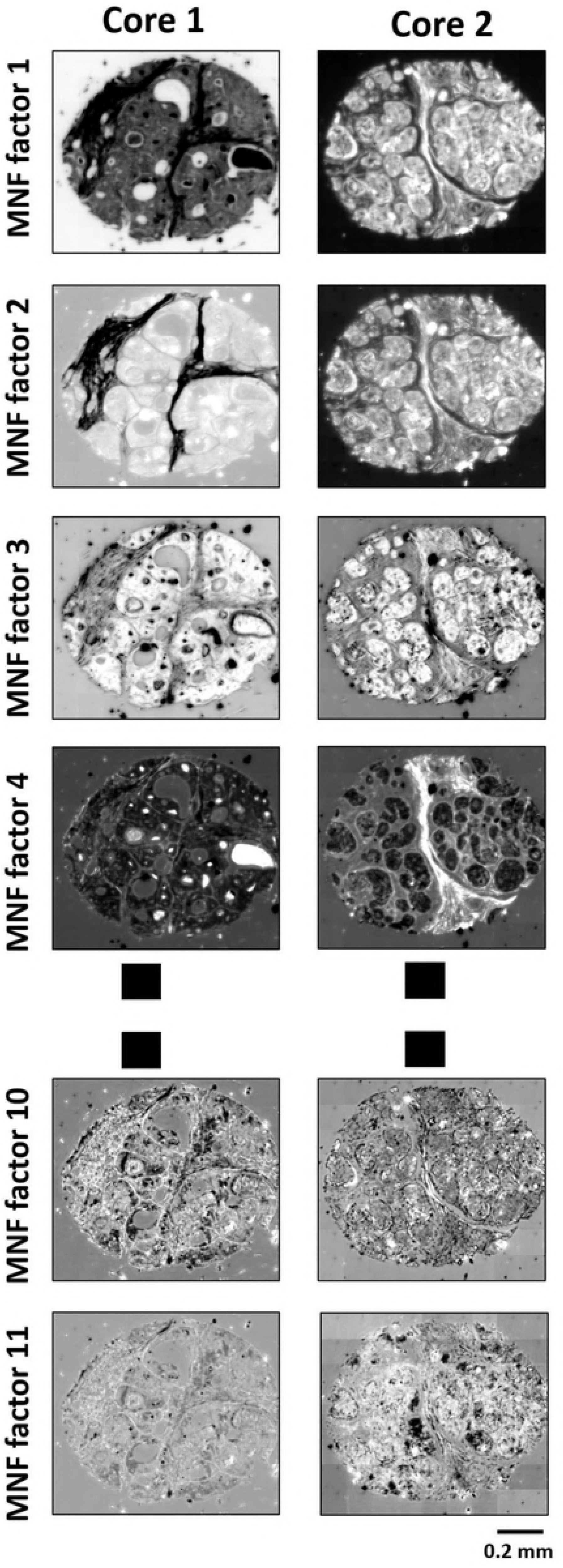
Spectral and SNR comparisons: A. IR image of a patient case at amide 1 band. B: Intensity profile along the horizontal line in the cases shown in A spanning the entire core for both the noisy and the denoised (in gray) version. C. Zoomed in view of the area marked with a red box in top row. D. Comparison of the spectral profile of the noisy and the MNF version at the pixel marked in red in C.

### Error Metric

Along with evaluating the performance of the presented MNF versions by examining the tissue profile, we utilize the MNI (method noise image) metric [24] aiming to maximize the structural similarity between the input noisy image and the denoised image around highly-structured regions, in the absence of ground truth. Fig. 3 shows the metric values for a core, over all the 1506 bands. A lower value of MNI indicate better denoising and structure preservation. This is evident from the fingerprint region (900 *−* 1800*cm^−1^*) which has very low values of MNI, while it is higher for the IR silent region (1800 *−* 2700*cm^−1^*). Since Standard MNF, Fast MNF and Approx MNF are almost the same in terms of reconstruction accuracy, all the plots from those three methods are same. The Rand MNF method which has a space-accuracy trade-off, produces similar results but with slight variations in most bands (notice the wavering in the plot).

**Fig 3.**
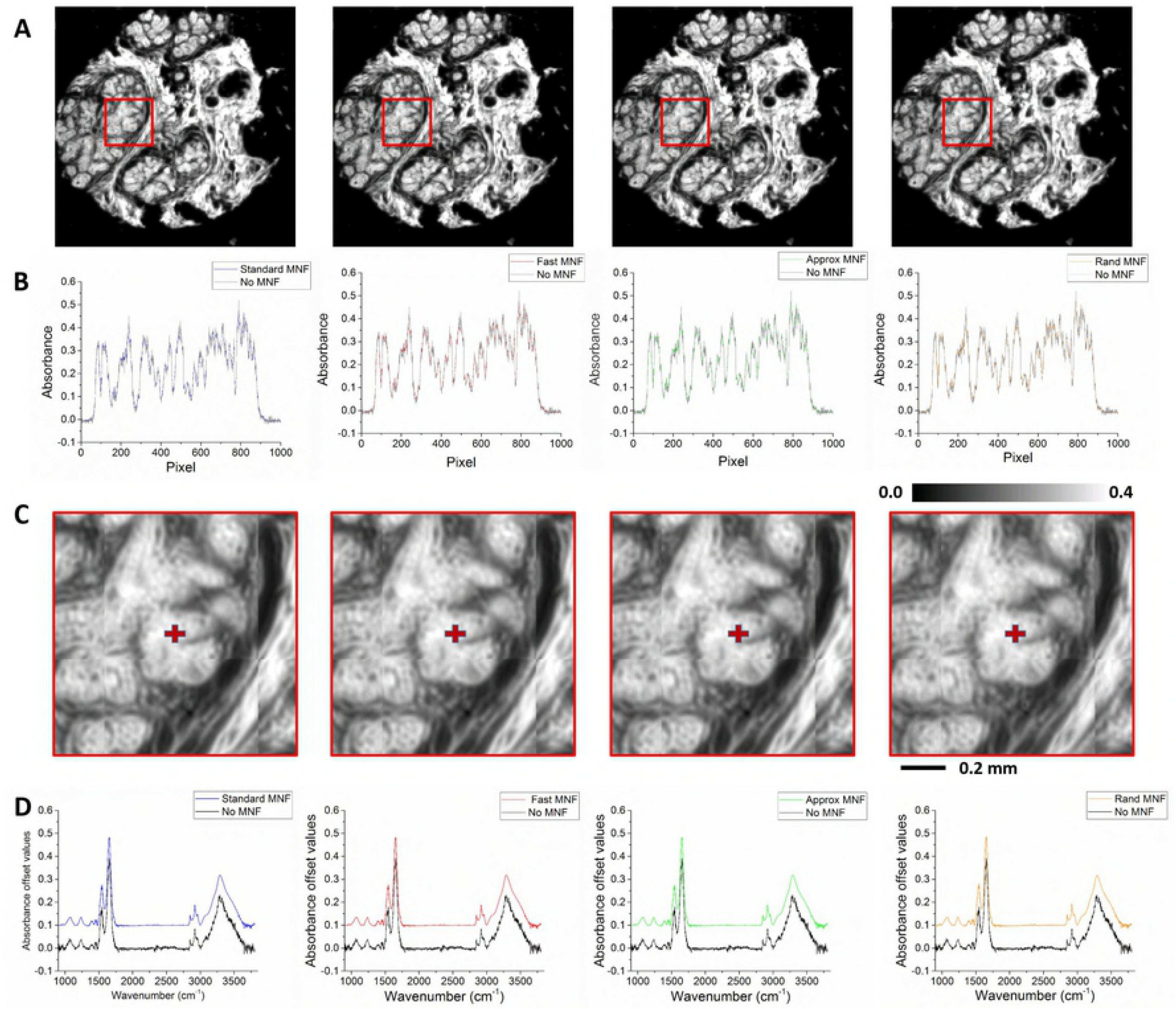
MNI metric plot for a breast tissue biopsy. Lower values of MNI indicate better denoising and perseverance of structure. Across all the four variations we notice low values of MNI in the fingerprint region (900 *−* 1800*cm^−1^*), which is mainly responsible for tissue classification. This shows that even in the presence of noise, the subtle structural features of the tissues are preserved after denoising. High MNI values in the IR silent region (1800 *−* 2700*cm^−1^*) are expected as they mainly contain noise and no relevant signal components. The functional region (2700 *−* 3600*cm^−1^*) has low MNI again followed by high MNI values in the water vapor absorption bands (3600 *−* 3800*cm^−1^*).

### Visualizing MNF bands

Fig. 4 visualizes the top *K* eigenimages (where *K* is automatically determined by the selection criteria) for two different patient cases (core1 and core 2). The bands in MNF space are arranged in decreasing order of SNR (increasing order of noise fraction), resulting in decreasing image quality with increase in band number. This suggests that the eigenimages in MNF space have decreasing image quality in terms of both the noise and structural detail. So,a few top bands in the MNF space should be able to capture most of information represented in the data along with denoising. This concept can further be utilized to develop automated selection criteria for the number of bands to be kept after the transform. This can help eliminate user based subjectivity, make the process faster and easier to implement.

**Fig 4.**
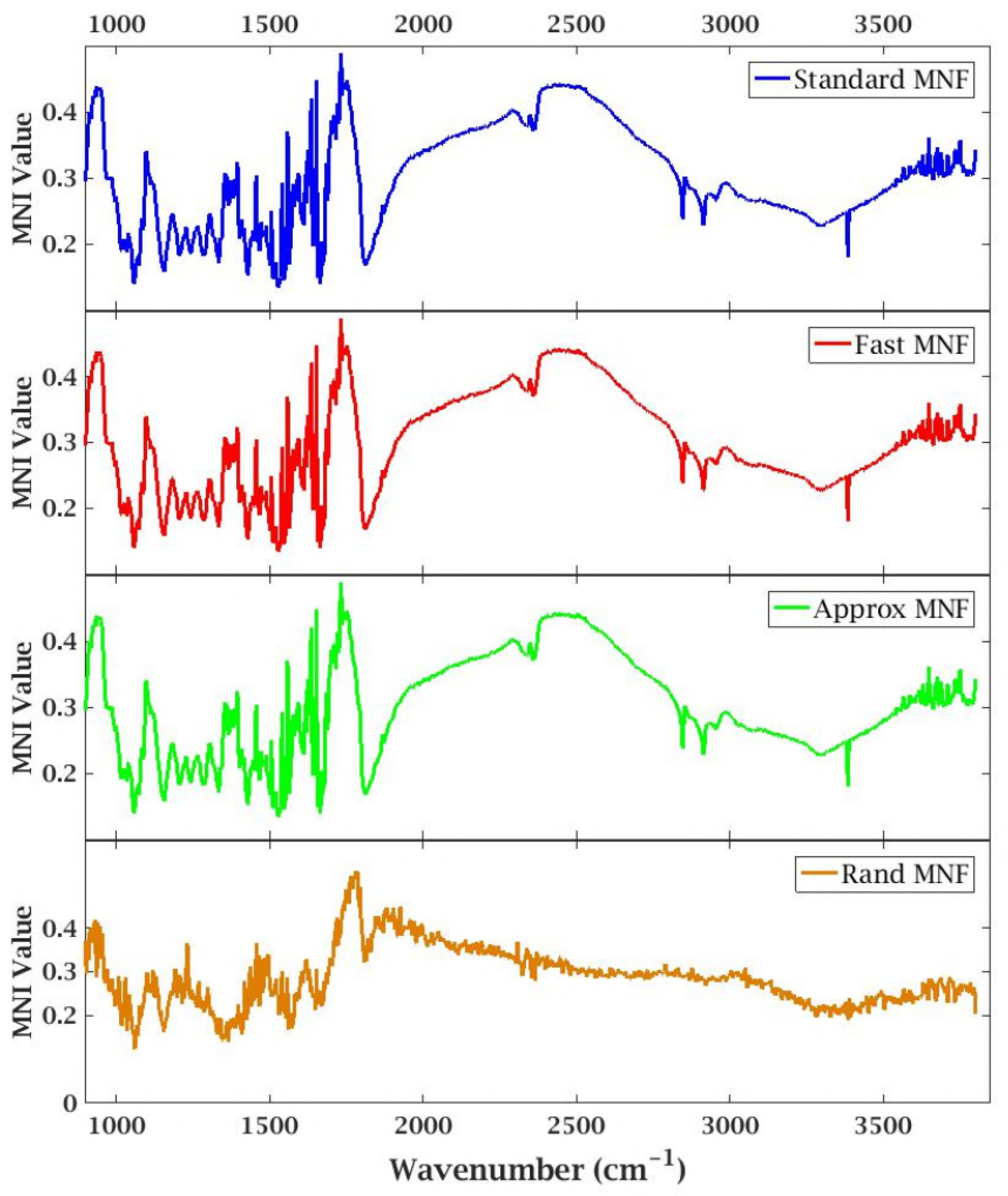
Eigenimages of the top K bands in the MNF space for two given TMA cores. There is an evident decrease in structural features and SNR with increase in band number. Manually inspecting these eigenimages or defining some measurement metric (Reddy *et al*. [19]) on them, increases the processing time and computation cost for MNF. Our approach automatically determines the optimal value of *K* from the MNF eigenvalues in a computationally efficient manner.

### Impact on Tissue Classification

Furthermore, we studied the effect of MNF based data processing on tissue classification models. In particular we have investigated the performance of different MNF algorithms against raw data for distinguishing cancer from benign breast cells. It can be seen in Fig. 5, that for all the MNF versions, the Area Under Curve (AUC) value of the malignant epithelium (cancer) class is the same and there is a 10% drop in accuracy without the use of MNF. This suggests that for the development of highly accurate and efficient diagnostic models with HSI data, MNF gives better performance. Also,the MNF techniques presented in this paper (Fast MNF, Rand MNF and Approx MNF), offers the same performance as compared to standard MNF with a factor of 60*×* reduction in the processing time and 50*×* reduction in memory space required for computation.

**Fig 5.**
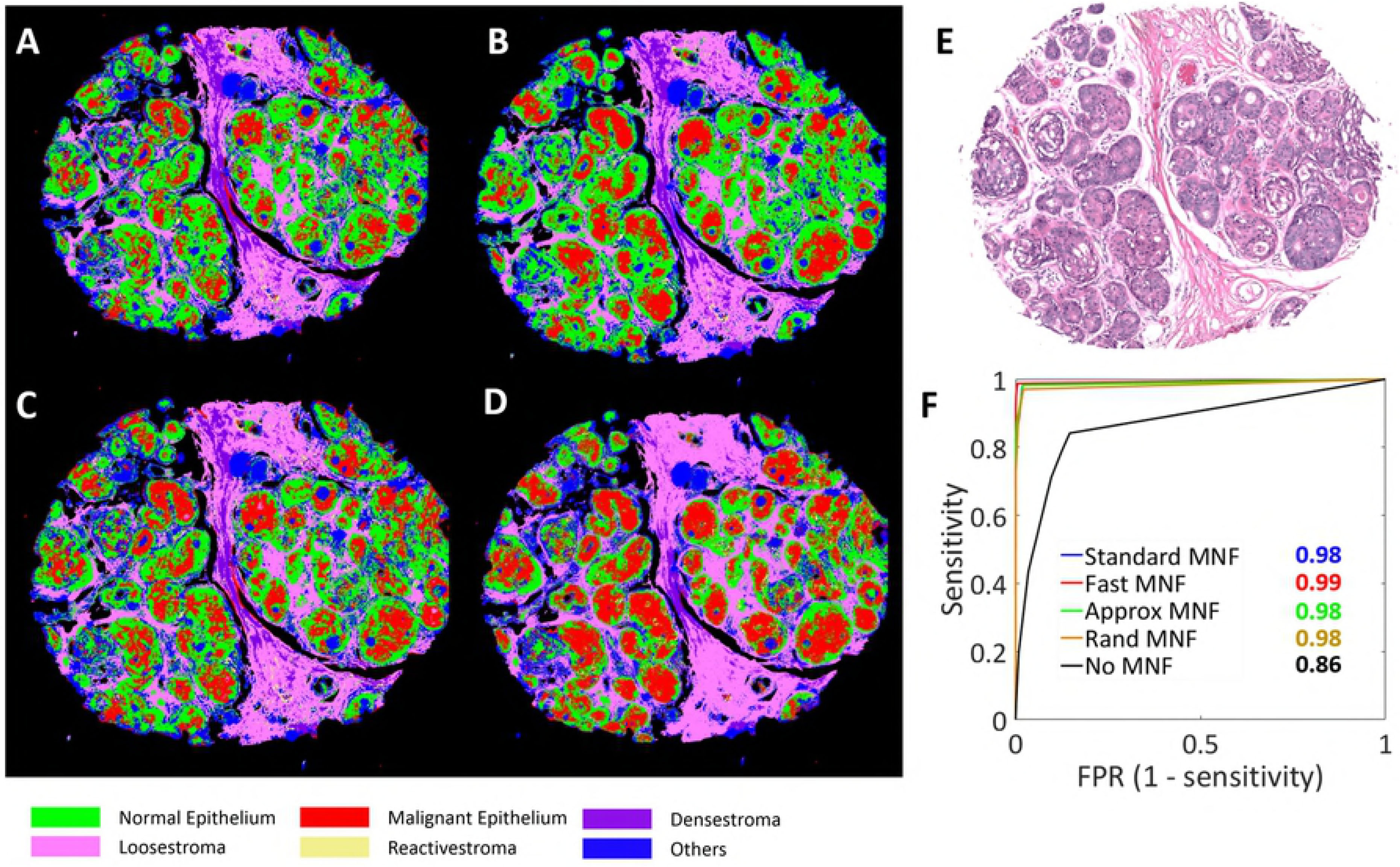
Effect of MNF variants on the classification accuracy of a breast tissue biopsy. A-D. Classified images after standard MNF, fast MNF, Approx MNF and Rand MNF in a clockwise order. E. H&E (Hematoxylin and Eosin) stained image of an adjacent slice of the same tissue. F. Receiver Operating Characteristic (ROC) curve with area under the curve (AUC) values for the malignant epithelium class.

## Conclusion

In this paper, we demonstrate how to automate the band selection process in the MNF space, which drastically reduces the workflow duration of MNF denoising of TMA’s from almost a month down to a matter of hours. We introduced three different optimizations of the MNF algorithm depending on the speed-memory-accuracy trade-off, resulting in a 60*×* runtime improvement and 50*×* memory efficiency. A well established error metric is also used which helps us decide the quality of denoising, in the absence of ground truth images. Similar classification performance of the suggested approaches as compared to conventional techniques suggesting the potential of the developed methods for computationally efficient analysis of big datasets for diagnostic applications. As a future work, we would like to make better approximations of the noise model itself, so that we can apply approximations for the eigen decomposition of the noise covariance matrix, hence further reducing the computational time of the process.

## Acknowledgments

This research was supported in part by an NIH grant R01GM117594.

